# Sift-PULs: A public repository for specific functional polysaccharide utilization loci

**DOI:** 10.1101/2021.08.04.455021

**Authors:** Tao Song, Congchong Wei, Dezhi Yuan, Shengwei Xiang, Hua Lv, Ting Huang, Kelei Zhao, Xinrong Wang, Yiwen Chu, Dan Luo, Jiafu Lin

**Author notes:** Corresponding authors of this paper **Corresponding author:** Dan Luo and Jiafu Lin, Sichuan Industrial Institute of Antibiotics, Chengdu University, Chengdu, 610106, P. R. China.

## Abstract

**Background:** Polysaccharide utilization loci (PULs) were bacterial gene clusters encoding genes responsible for polysaccharide utilization process. PUL studies are blooming in recent years but the biochemical characterization speed is relative slow. There is a growing demand for PUL database with function annotations.

**Results:** Using signature genes corresponding for specific polysaccharide, 10422 PULs specific for 6 polysaccharides (agar, alginate, pectin, carrageenan, chitin and β-manan) from various bacterial phyla were predicted. Then online website of specific functional polysaccharide utilization loci (Sift-PULs) was constructed. Sift-PULs provides a repository where users could browse, search and download interested PULs without registration.

**Conclusions:** The key advantage of Sift-PULs is to assign a function annotation of each PUL, which is not available in existing PUL databases. PUL’s functional annotation lays a foundation for studying novel enzymes, new pathways, PUL evolution or bioengineering. The website is available on https://sift-puls.org

## Introduction

PUL s(polysaccharide utilization loci) are bacterial gene clusters that encoding a variety of functional genes responsible for the transcription, degradation, transport and metabolism of polysaccharides (Grondin *et al.*, 2017). Since the discovery of starch PUL (also named as Sus. starch utilization system) from *Bacteroides thetaiotaomicron*, more and more PULs have been found from different ecosystems and various bacteria phyla (D’Elia and Salyers, 1996; Foley *et al.*, 2016; Chen *et al.*, 2018; Despres *et al.*, 2016; Grondin *et al.*, 2017). Discovered PULs are found to target at many different types of polysaccharides, including xylan, β-mannan or pectin (Tang *et al.*, 2017; Bagenholm *et al.*, 2017; Reddy *et al.*, 2016; Ficko-Blean *et al.*, 2017; Despres *et al.*, 2016; Pluvinage *et al.*, 2018). Now PUL studies are becoming a hotspot because their significant importance in ecology, evolution and bioengineering(Grondin *et al.*, 2017). Although PULs have important biological functions, biochemical identification in laboratory is too slow, resulting in number scarcity and hindering the progress of PUL study.

In view of the sparse data of PULs and PULs’ significant biological functions, it is very necessary to use bioinformatics way to identify PULs. There are currently three PUL databases available, including PULDB, CGCs and dbCAN-PUL(Terrapon *et al.*, 2015; Zhang *et al.*, 2018; Ausland *et al.*, 2021). PULDB is the first PUL database, which uses SusC/D gene pair and carbohydrate active enzymes for prediction. It mainly includes PULs from *Bacteroidetes*. CGCs predict PULs using transcription factors, transport proteins and carbohydrate active enzymes while dbCAN-PUL does not predict PULs but provides experimentally confirmed PULs. It is worth mentioning that although these three databases provide PUL collections, there is no annotation for predicted PULs. So researchers who want to find PULs targeting at specific polysaccharide need to manually check the predicted PULs. This is very laborious. PUL with a function annotation is helpful for researchers to answer new hypotheses and provide a basis for new discoveries. With the increase of PUL studies, the requirement of PUL database with specific function annotations has become more and more urgent. Unfortunately, there is no PUL database providing specific function annotation now.

PULDB and CGCs are the two main databases currently used for PUL prediction, and no functional prediction is given for the predicted PULs. Therefore, we used signature genes corresponding to 6 different polysaccharides (agar, alginate, pectin, carrageenan, chitin and β-manan) to predict PULs, and gave function predictions. Then Sift-PULs website is constructed, where users could easily search and download interested Sift-PULs. Sift-PULs serves as repository for researchers who focus on one specific polysaccharide and need large scale data to discover novel protein, utilization pathways or evolutional process.

## Materials and methods

### Data retrieval

Bacterial genomes were mainly downloaded from NCBI database (download was finished in 2021.03.01). Genomes at different assemble levels (contig, scaffold, complete or chromosome) were downloaded using a home-made script. Only GBFF format file were retrieved from FTP link.

### Data normalization

The genbank file of a bacterial genome was parsed using Biopython package, then protein sequences within bacterial genome were extracted into a single fasta file. The name of each protein is normalized into following format: GCF number of genome, contig name, serial number on the contig, gene start position, gene end position and gene direction. Therefore, protein name was a unique signature which contained essential information for prediction.

### Selection of signature gene

In total, PULs that were specific for alginate, agar, carrageenan, chitin, β-mannan and pectin (polygalacturonicacid) were considered in this manuscript. In this study, signature genes were classified into two categories, core genes and alternative genes (Supplementary material 1). Core genes referred to genes that were essential for the polysaccharide utilization process, including ones responsible for monosaccharide metabolism (e.g. unique 3,6 anhydro-L galactose metabolic genes for agar) or unique metabolic process (e.g. GH130 mannobiose phosphorylase for mannan utilization). Core genes were determined if they were commonly found in most biochemically PULs. Alternative genes were usually carbohydrate active enzymes responsible for polysaccharide utilization. Alternative genes were determined if they appeared in characterized PULs or their activities were related to polysaccharide degradation.

### Hmmer model build

Most hmmer models responsible for signature genes were built locally. To build an hmmer model, experimentally validated protein sequences were first collected and aligned using MUSCLE(Edgar, 2004), followed by manual correction. Proteins with experimental evidences from CAZYs and Unipro were used as test data to test the true positive rate and false positive rate. The re-build hmmer model should have >95% true positive rate and <5% false positive rate under a specific threshold. When the signature gene was from a family with only one enzyme activity and this family had very few experimentally confirmed members (less than 3), the corresponding hmmer model was retrieved from Pfam and dbCAN (Finn *et al.*, 2013; Zhang *et al.*, 2018).

### Sift-PULs prediction

Sift-PUL prediction needed 6 steps:

1. Firstly, normalized protein fasta file was analyzed using Hmmer against corresponding models. If domain of a gene was the same as core gene or alternative gene, it was recorded as core gene or alternative gene, respectively.
2. Then, we investigated whether it was possible to put core genes and at least one alternative gene into a gene cluster with less than 50 genes. If did, serial numbers of matched core genes or alternative genes were recorded.
3. Minimum PUL was defined as gene cluster contained minimum members including all core genes and at least one alternative gene. Extended the minimum PUL to both sides until the gene number reached 50. Then extended PUL was defined as maximum PUL.
4. Calculate the frequency of individual domain in minimum PULs. Domains with >10% frequency were defined as high frequency domain.
5. PUL boundary of minimum PUL was extended until the adjacent and consecutive 5 genes did not have high frequency domain. If the extended PULs were smaller than maximum PUL, extended boundary was used. Otherwise, maximum PUL boundary was used.

### Online database construct

The website of Sift-PULs was constructed using Vue.js (javascript) and Django (python). Database is implemented using PosgreSQL.

## Result and discussion

### Data collections of Sift-PULs

PULs are bacterial gene clusters that have essential biological functions. Considering increasing interest of researchers in PUL study and PULs’ slow identification in laboratory, it is necessary to establish a PUL database with function prediction. Using signature genes that specific to corresponding polysaccharide, 10422 PULs were identified, including 2347 pectin PULs, 1140 manan PULs, 1938 alginate PULs, 4723 chitin PULs, 186 agar PULs and 88 carrageenan PULs (Fig 1A). Meanwhile, predicted PULs came from different phyla including *Proteobacteria* (4140 PULs), *Firmicutes* (3335 PULs), *Bacteroidetes* (2342 PULs) and *Actinobacteria* (537 PULs). Noteworthy, Sift-PULs showed a potent as a reference database for discovering novel PULs. For example, predicted carrageenan PULs came from 5 phyla (*Actinobacteria*, *Bacteroidetes*, *Planctomycetes*, *Proteobacteria* and *Firmicutes*), and now only carrageenan from *Bacteroidetes* was experimentally verified. Moreover, bacterial genomes above contig level were used for sift-PULs prediction in this study, therefore most predicted sift-PULs come from bacteria had contig or scaffold genomes (Fig 1B).

**Figure 1.**
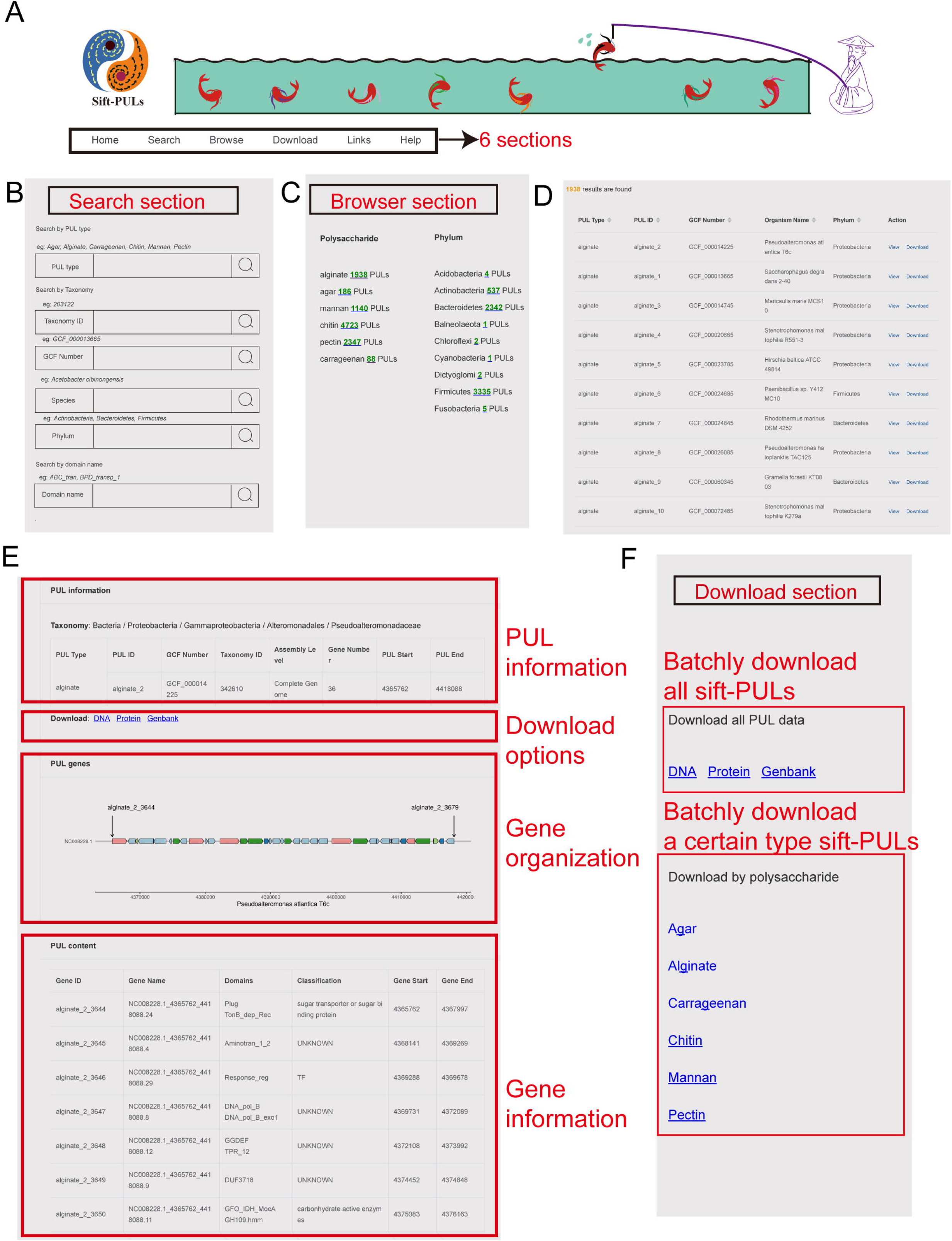
summary of predicted sift-PULs. A: phyla distribution of bacteria with predicted sift-PULs. B: Assemble level of bacteria with predicted sift-PULs.

**Figure 2:**
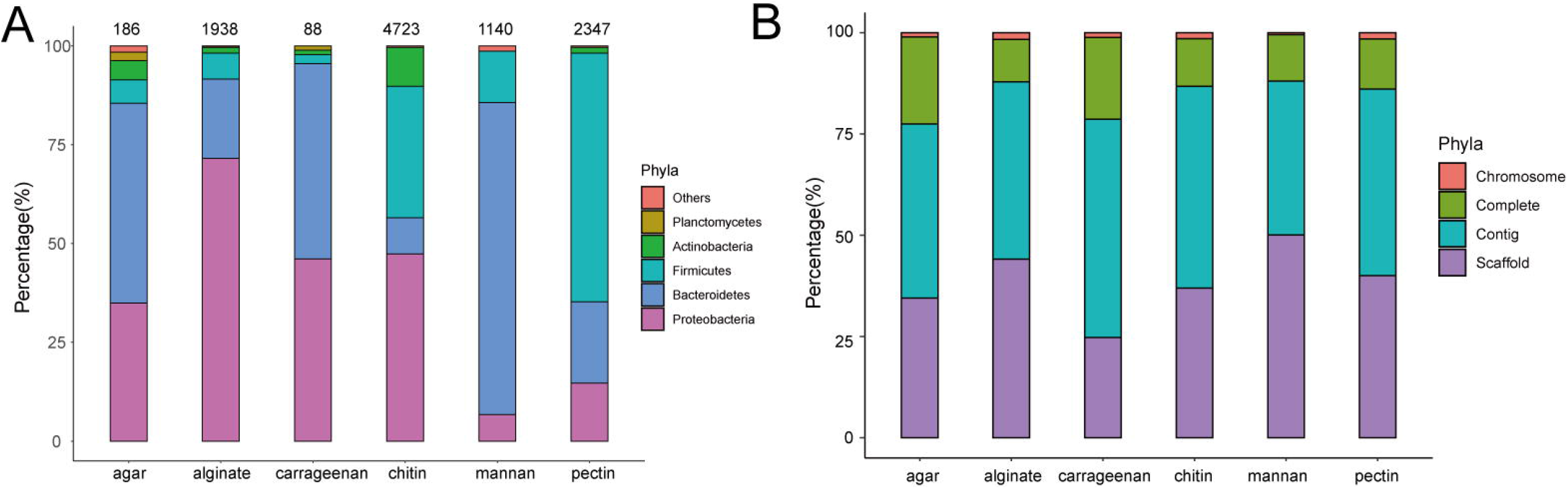
Screenshots of sift-PULs website. A: Menu in sift-PULs website link to different sections. B: Search section in sift-PULs website. Users could search sift-PULs by phyla, species name, txa id and domain name. C: Browse section in sift-PULs website. Sift-PULs were classified by polysaccharide or phyla. D: Screenshot of sift-PULs list after user click search or browse button. E: Web interface for a sift-PUL using a alginate PUL as an example. F: Web interface for batch download.

In current study, hmmer models were locally built to ensure signature genes’ specificity, then combination of signature genes was used for PUL’s function prediction. Still, it was possible that our predicted results may contain false positives.

To investigate the data reliability of predicted sift-PULs, firstly, we tried to evaluate the prediction method to give hint about data accuracy. However, it did not succeed because of insufficient number of experimentally confirmed PULs. For example, there was only one report of carrageenan PUL, 2 reports of agar PULs, less than 10 reports of pectin PULs. Then, we tried to find evidence in database containing bacterial polysaccharide utilization information (biodive). However, bacteria with sift-PULs were either not recorded in biodive database, or corresponding polysaccharide utilization information was not recorded. Matched results were too few to get any useful conclusions. In the end, we focused on bacteria with agar-PULs. This was because agar was commonly used in bacterial cultivation for almost 100 years, agar degradation phenotype could be easily seen on plate, and this information was more likely to be recorded in literature. Surprisingly, 65 out of 186 bacteria with predicted agar PULs could degrade agar and the rest were not mentioned (Supplementary material 2). This implied the predicted agar PULs were relative reliable. The accuracy of agar PULs also indicated sift-PULs could be used as reference databank for researchers.

### Comparison with existing PUL databases

Using signature genes to predict PUL was commonly in current research(Terrapon *et al.*, 2015; Zhang *et al.*, 2018). For example, SIFT-PULS used the PUL conservative SusC/D gene pair and carbohydrate active enzymes, and CGCs used transcription factors, sugar transporters and carbohydrate active enzymes. The signature genes used in prediction determine the properties of the obtained PUL. For example, the PULs in SIFT-PULS only came from *Bacteroidetes*, because the SusC/D gene pair was mainly derived from *Bacteroidetes*. Because the selected signature genes are not specific to polysaccharides in PULDB or CGCs, none of these two databases could give function prediction. The signature genes used in Sift-PULs in this article were specific to each polysaccharide, therefore function annotation was possible. The function prediction greatly reduced the workload of researchers searching for the corresponding function PUL.

Sift-PULs included 10422 PULs, which less than with 43156 PULs in PULDB (Table 1). This was probably because sift-PULs only focus on 6 polysaccharides. Meanwhile, PULs from Sift –PULS were from multiple bacterial phyla. This was similar to CGCs but different with sift-PULS, in which only *Bacteroidetes* was considered. Compared with sift-PULs and CGCs, the most important feature of Sift-PULs was that it could give function predictions.

**Table 1.** Comparison Sift-PULDB with PULDB and CGCs, N.A.: not available

|  | PULDB | CGCs | Sift-PULs |
| --- | --- | --- | --- |
| Signature Genes | SusC/D and CAZys | Transcription factor, transporter and CAZys | Specific genes for each polysaccharide |
| PUL numbers | 43156 | N.A. | 10422 |
| Multiple phyla | No | Yes | Yes |
| Bacteria assemble level | Complete | Complete | above contig |
| Function prediction | No | No | yes |

### Web interface

At the start page of sift-PULs, there were six sections: home, search, browse, download, links and help (Fig 1A). At home page, there was a brief introduction of sift-PULs, where users could quickly learn how sift-PULs were predicted. Important update would also be showed here. User could find interested sift-PULs in two ways. First, in the search section, users could search for interested PULs using different keywords, for example polysaccharide name, taxid, GCF number, phylum name, species name or protein domain name (Fig 1B). Second, in browse section, sift-PULs were classified by polysaccharide or phyla (Fig 1C). By clicking the search button in search section or links in browse section, interested sift-PULs would be displayed (Fig 1D). After clicking ‘view’ button interested PUL, PUL information would be displayed in a pop-up page, which contains the PUL information, download option, gene cluster map and gene information(Fig 1E)..

Sift-PULs also provide batch download service, which was convenient for users who required large amount of data. In download section, users could easily download all sift-PULs data (Fig 1F). There were three format files available, including a genbank file that included complete DNA sequence of PUL, DNA fasta file that included DNA sequences for individual CDS, protein fasta file that included protein sequences for individual CDS. Users could download these files when browsing individual PUL.

### Conclusions

Sift-PULs website provides a public repository where users could easily access, search and download PULs with specific function annotation, which helps researchers to build a local database and come up with novel hypothesis. For example, Sift-PULs could help biochemists discover novel enzymes (study proteins that are not characterized but have high frequency score) and find novel degradation pathways. In future, Sift-PULs would update once a year. Update would include sift-PULs from newly sequenced genomes or sift-PULs targeting at new polysaccharides (e.g. α-mannan, starch and ulvan).

Online prediction service of sift-PULs is also under construction.

## Declaration

### Abbreviations (if applicable)

PULs: polysaccharide utilization loci

### Ethics approval and consent to participate

Not applicable

### Consent for publication

Not applicable

### Availability of data and materials

Not applicable

## Acknowledgements

This work was supported by the Sichuan Science and Technology Program (Grant Number:2019YJ0282); National Natural Science Foundation of China (31970131),Collaboration and Innovation on New

Anti-biotic Development and Industrialization (Grant Number: 2016-XT00-00023-G); Major New Drugs

Innovation and Development (Grant Number: 2018ZX09711-001); The talent Introduction of Project of Chengdu University (2081918021) and Key Laboratory of Coarse Cereal Processing, Ministry of Agriculture and Rural Affairs (2020CC009). Publishing cost was funded by Antibiotics Research and Re-evaluation Key Laboratory of Sichuan Province (Grant Number: ARRLKF19-01, ARRLKF19-05); The Science and technology projects from Department of Ecology and Environment of Sichuan Province (Grant No. 2019HB16). This work was supported by Enzyme Resources Sharing and Service Platform of Sichuan Province.-Competing Interests

We declare that we have no competing interest.

Acknowledgements

We thank Dr. Jan Hendrik Hehemann (Max Planck Institute for Marine Microbiology) for his critical comments on polysaccharide utilization pathways.

