## Supplementary material for "Sift-PULs: A public repository for specific functional polysaccharide utilization loci"

Supplementary material 1: Information summary of selected core genes and alternative genes. In Activity column, MME represent monosaccharide metabolic enzymes, a: enzyme family from Pfam database, b: enzyme family from CAZY database. c: cut value that used for corresponding model.

| polysaccharide | activity | Classification | Enzyme family | Rebuild model? | cut_value[c] |
|---|---|---|---|---|---|
| Agar | MME | Core gene | PF00171[a] | Y | 1.00E-100 |
| | MME | Core gene | PF13378[a] | Y | 1.00E-50 |
| | MME | Core gene | PF08240[a] | Y | 1.00E-50 |
| | MME | Core gene | PF 00106[a] | Y | 1.00E-50 |
| | agarase | Alternative gene | GH16[b] | Y | 1.00E-25 |
| | agarase | Alternative gene | GH86[b] | Y | 1.00E-20 |
| | agarase | Alternative gene | GH117[b] | Y | 1.00E-20 |
| | galactosidase | Alternative gene | GH2[b] | Y | 1.00E-50 |
| | agarase | Alternative gene | GH96[b] | Y | 1.00E-10 |
| | agarase | Alternative gene | GH118[b] | Y | 1.00E-10 |
| carrageenan | MME | Core gene | PF00171[a] | Y | 1.00E-50 |
| | MME | Core gene | PF13378[a] | Y | 1.00E-50 |
| | carrageenase | Alternative gene | GH82[b] | Y | 1.00E-20 |
| | carrageenase | Alternative gene | GH16[b] | Y | 1.00E-50 |
| | carrageenase | Alternative gene | GH150[b] | Y | 1.00E-50 |
| pectin | MME | Core gene | PF02614[a] | N | 1.00E-10 |
| | MME | Core gene | PF04295[a] | N | 1.00E-10 |
| | MME | Core gene | PF08125[a] | N | 1.00E-10 |
| | pectatelyase | Alternative gene | PL1[b] | Y | 1.00E-10 |
| | pectatelyase | Alternative gene | PL10[b] | Y | 1.00E-10 |
| | pectatelyase | Alternative gene | PL9[b] | Y | 1.00E-50 |
| | pectatelyase | Alternative gene | PL3[b] | Y | 1.00E-10 |
| | pectatelyase | Alternative gene | PL2[b] | Y | 1.00E-10 |
| | pectinmethylesterase | Alternative gene | CE8[b] | Y | 1.00E-10 |
| mannan | MME | Core gene | PF01182[a] | N | 1.00E-10 |
| | mannanase | Core gene | GH130[b] | N | 1.00E-10 |
| | mannanase | Alternative gene | GH5[b] | Y | 1.00E-25 |
| | mannanase | Alternative gene | GH26[b] | Y | 1.00E-20 |
| | mannosidase | Alternative gene | GH2[b] | Y | 1.00E-50 |
| | mannosidase | Alternative gene | GH5[b] | Y | 1.00E-50 |
| | mannanase | Alternative gene | GH134[b] | N | 1.00E-10 |
| | mannanase | Alternative gene | GH164[b] | N | 1.00E-10 |
| chitin | MME | Core gene | PF07221[a] | N | 1.00E-10 |
| | Esterase | Core gene | CE9[b] | N | 1.00E-10 |
| | chitinase | Alternative gene | GH18[b] | Y | 1.00E-10 |
| | chitinase | Alternative gene | GH19[b] | Y | 1.00E-10 |
| | chitinase | Alternative gene | GH75[b] | N | 1.00E-10 |
| | chitinase | Alternative gene | GH80[b] | N | 1.00E-10 |
| | acetylglucosaminidase | Alternative gene | GH18[b] | Y | 1.00E-30 |

| | | | | | |
|---|---|---|---|---|---|
| | acetylglucosaminidase | Alternative gene | GH73 [b] | Y | 1.00E-10 |
| | acetylglucosaminidase | Alternative gene | GH84 [b] | Y | 1.00E-10 |
| | acetylglucosaminidase | Alternative gene | GH20 [b] | Y | 1.00E-10 |
| | acetylglucosaminidase | Alternative gene | GH3 [b] | Y | 1.00E-50 |
| | chitosanase | Alternative gene | GH46 [b] | Y | 1.00E-10 |
| | chitosanase | Alternative gene | GH8 [b] | Y | 1.00E-150 |
| alginate | MME | Core gene | PF13561 [a] | Y | 1.00E-40 |
| | alginatelyase | Alternative gene | PL7 [b] | Y | 1.00E-10 |
| | alginatelyase | Alternative gene | PL6 [b] | Y | 1.00E-10 |
| | alginatelyase | Alternative gene | PL5 [b] | Y | 1.00E-10 |
| | alginatelyase | Alternative gene | PL31 [b] | Y | 1.00E-50 |
| | alginatelyase | Alternative gene | PL17 [b] | N | 1.00E-10 |
| | alginatelyase | Alternative gene | PL34 [b] | N | 1.00E-10 |
| | alginatelyase | Alternative gene | PL15 [b] | Y | 1.00E-10 |

Supplementary material 2：Information of bacterial genomes that had predictedagar PUL

| Genome name | Phylum | GCA-number |
|---|---|---|
| *Streptomyces coelicolor* A3(2)[1] | *Actinobacteria* | GCA_000203835.1 |
| *Sphingomonas* sp. MCT13 [2] | *Alphaproteobacteria* | GCA_001721295.1 |
| *Algibacter lectus*[3] | *Bacteroidetes/Chlorobi group* | GCA_000764755.1 |
| *Algibacter lectus*[3] | *Bacteroidetes/Chlorobi group* | GCA_900112395.1 |
| *Aquimarina agarilytica* ZC1[4] | *Bacteroidetes/Chlorobi group* | GCA_000255455.1 |
| *Aquimarina latercula* DSM 2041[5] | *Bacteroidetes/Chlorobi group* | GCA_000430645.1 |
| *Aquimarina* sp. RZW4-3-2[6] | *Bacteroidetes/Chlorobi group* | GCA_001632745.1 |
| *Cellulophaga algicola* DSM 14237[7] | *Bacteroidetes/Chlorobi group* | GCA_000186265.1 |
| *Cellulophaga baltica*[8] | *Bacteroidetes/Chlorobi group* | GCA_900102165.1 |
| *Cellulophaga baltica* NN016038[8] | *Bacteroidetes/Chlorobi group* | GCA_000477035.2 |
| *Cellulophaga fucicola*[8] | *Bacteroidetes/Chlorobi group* | GCA_900119145.1 |
| *Cellulophaga lytica*[8] | *Bacteroidetes/Chlorobi group* | GCA_000750195.1 |
| *Cellulophaga lytica*[8] | *Bacteroidetes/Chlorobi group* | GCA_001941605.1 |
| *Cellulophaga lytica* DSM 7489 [9] | *Bacteroidetes/Chlorobi group* | GCA_000190595.1 |

| | | |
|---|---|---|
| *Cellulophaga* sp. E6(2014)[10] | *Bacteroidetes/Chlorobi group* | GCA_000764435.1 |
| *Echinicola pacifica* DSM 19836[11] | *Bacteroidetes/Chlorobi group* | GCA_000373245.1 |
| *Flammeovirga pacifica*[12] | *Bacteroidetes/Chlorobi group* | GCA_000807855.2 |
| *Flammeovirga* sp. MY04[13] | *Bacteroidetes/Chlorobi group* | GCA_001682195.1 |
| *Flavobacterium flevense*[14] | *Bacteroidetes/Chlorobi group* | GCA_900142775.1 |
| *Formosa agariphila* KMM 39011[5] | *Bacteroidetes/Chlorobi group* | GCA_000723205.1 |
| *Formosa haliotis*[16] | *Bacteroidetes/Chlorobi group* | GCA_001685485.1 |
| *Maribacter aquivivus*[17] | *Bacteroidetes/Chlorobi group* | GCA_900142175.1 |
| *Ochrovirga pacifica*[18] | *Bacteroidetes/Chlorobi group* | GCA_000220525.2 |
| *Persicobacter* sp. JZB09[19] | *Bacteroidetes/Chlorobi group* | GCA_001308105.1 |
| *Polaribacter reichenbachii*[20] | *Bacteroidetes/Chlorobi group* | GCA_001975665.1 |
| *Polaribacter reichenbachii*[20] | *Bacteroidetes/Chlorobi group* | GCA_001680875.1 |
| *Pseudozobellia thermophila*[21] | *Bacteroidetes/Chlorobi group* | GCA_900141855.1 |
| *Reichenbachiella agariperforans*[22] | *Bacteroidetes/Chlorobi group* | GCA_900142205.1 |
| *Saccharicrinis fermentans* DSM 9555 = JCM 21142[2324] | *Bacteroidetes/Chlorobi group* | GCA_000517085.1 |
| *Tamlana agarivorans*[25] | *Bacteroidetes/Chlorobi group* | GCA_001642835.1 |
| *Tamlana* sp. s12 [26] | *Bacteroidetes/Chlorobi group* | GCA_001672305.1 |
| *Zobellia galactanivorans*[27] | *Bacteroidetes/Chlorobi group* | GCA_000973105.1 |
| *Zobellia uliginosa*[28] | *Bacteroidetes/Chlorobi group* | GCA_000744555.1 |
| *Wenyingzhuangia fucanilytica*[29] | *Bacteroidetes/Chlorobi group* | GCA_ 001697185.1 |
| *Aquimarina aggregata*[6] | *Bacteroidetes/Chlorobi group* | GCA_ 001632745.1 |
| *Echinicola strongylocentroti*[30] | *Bacteroidetes/Chlorobi group* | GCA_ 003260975.1 |

| | | |
|---|---|---|
| *Labilibacter* sp. CG51[31] | *Bacteroidetes/Chlorobi group* | GCA_ 005877885.1 |
| *Reichenbachiella versicolor*[32] | *Bacteroidetes/Chlorobi group* | GCA_ 003171675.1 |
| *Sediminitomix flava*[33] | *Bacteroidetes/Chlorobi group* | GCA_003149185.1 |
| *Algoriphagus chordae*[34] | *Bacteroidetes/Chlorobi group* | GCA_ 003254055.1 |
| *Cellvibrio* sp. OA-2007[35] | *Gammaproteobacteria* | GCA_000953825.1 |
| *Cellvibrio* sp. Pealriver[36] | *Gammaproteobacteria* | GCA_001183545.1 |
| *Gayadomonas joobiniege* G7[37] | *Gammaproteobacteria* | GCA_000300815.1 |
| *Gilvimarinus agarilyticus*[38] | *Gammaproteobacteria* | GCA_000832015.1 |
| *Gilvimarinus chinensis* DSM 19667[39] | *Gammaproteobacteria* | GCA_000377745.1 |
| *Gilvimarinus polysaccharolyticus*[40] | *Gammaproteobacteria* | GCA_001187555.1 |
| *Marinimicrobium agarilyticum* DSM 16975 [41] | *Gammaproteobacteria* | GCA_000423345.1 |
| *Microbulbifer agarilyticus* S89 [42] | *Gammaproteobacteria* | GCA_000220505.2 |
| *Microbulbifer* sp. HZ11[43] | *Gammaproteobacteria* | GCA_000708675.1 |
| *Microbulbifer thermotolerans*[44] | *Gammaproteobacteria* | GCA_001617625.1 |
| *Microbulbifer thermotolerans*[45] | *Gammaproteobacteria* | GCA_900112305.1 |
| *Paraglaciecola* sp. S66[46] | *Gammaproteobacteria* | GCA_001565895.1 |
| *Pseudoalteromonas mariniglutinosa*[47] | *Gammaproteobacteria* | GCA_001662245.1 |
| *Saccharophagus degradans* 2-40 [48] | *Gammaproteobacteria* | GCA_000013665.1 |
| *Simiduia agarivorans*[49] | *Gammaproteobacteria* | GCA_000305785.2 |
| *Simiduia agarivorans*[49] | *Gammaproteobacteria* | GCA_000420285.1 |
| *Vibrio* sp. EJY3 [50] | *Gammaproteobacteria* | GCA_000241385.1 |
| *Shewanella pacifica*[51] | *Gammaproteobacteria* | GCA_ 002075795.1 |
| *Steroidobacter agariperforans*[52] | *Gammaproteobacteria* | GCA_ 004138335.1 |
| *Agaribacterium haliotis*[53] | *Gammaproteobacteria* | GCA_002312815 |
| *Alteromonadaceae bacterium* ALS 81[54] | *Gammaproteobacteria* | GCA_ 003610775.1 |


1. M. J. Buttner, I. M. Fearnley, M. J. Bibb, The agarase gene (dagA) of Streptomyces coelicolor A3(2): nucleotide sequence and transcriptional analysis. *Mol. Gen. Genet. MGG*. **209**, 101–109 (1987).
2. M. M. D'Andrea, N. Ciacci, V. Di Pilato, G. M. Rossolini, M. C. Thaller, Draft Genome Sequence of the Agarase-Producing Sphingomonas sp. MCT13. *Front. Environ. Sci.***5** (2017), doi:10.3389/fenvs.2017.00009.

3. O. I. Nedashkovskaya, S. B. Kim, S. K. Han, M.-S. Rhee, A. M. Lysenko, M. Rohde, N. V. Zhukova, G. M. Frolova, V. V. Mikhailov, K. S. Bae, Algibacter lectus gen. nov., sp. nov., a novel member of the family Flavobacteriaceae isolated from green algae. *Int. J. Syst. Evol. Microbiol.***54**, 1257–1261 (2004).
4. B. Lin, G. Lu, S. Li, Z. Hu, H. Chen, Draft Genome Sequence of the Novel Agarolytic Bacterium Aquimarina agarilytica ZC1. *J. Bacteriol.***194**, 2769–2769 (2012).
5. O. I. Nedashkovskaya, M. Vancanneyt, L. Christiaens, N. I. Kalinovskaya, V. V. Mikhailov, J. Swings, Aquimarina intermedia sp. nov., reclassification of Stanierella latercula (Lewin 1969) as Aquimarina latercula comb. nov. and Gaetbulimicrobium brevivitae Yoon et al. 2006 as Aquimarina brevivitae comb. nov. and emended description of the genus Aquimarina. *Int. J. Syst. Evol. Microbiol.***56**, 2037–2041 (2006).
6. Y. Wang, H. Ming, W. Guo, H. Chen, C. Zhou, Aquimarina aggregata sp. nov., isolated from seawater. *Int. J. Syst. Evol. Microbiol.***66**, 3406–3412 (2016).
7. J. P. Bowman, Description of Cellulophaga algicola sp. nov., isolated from the surfaces of Antarctic algae, and reclassification of Cytophaga uliginosa (ZoBell and Upham 1944) Reichenbach 1989 as Cellulophaga uliginosa comb. nov. *Int. J. Syst. Evol. Microbiol.***50**, 1861–1868 (2000).
8. J. E. Johansen, P. Nielsen, C. Sjøholm, Description of Cellulophaga baltica gen. nov., sp. nov. and Cellulophaga fucicola gen. nov., sp. nov. and reclassification of [Cytophaga] lytica to Cellulophaga lytica gen. nov., comb. nov. *Int. J. Syst. Evol. Microbiol.***49**, 1231–1240 (1999).
9. B. Abt, M. Lu, M. Misra, C. Han, M. Nolan, S. Lucas, N. Hammon, S. Deshpande, J.-F. Cheng, R. Tapia, L. Goodwin, S. Pitluck, K. Liolios, I. Pagani, N. Ivanova, K. Mavromatis, G. Ovchinikova, A. Pati, A. Chen, K. Palaniappan, M. Land, L. Hauser, Y.-J. Chang, C. D. Jeffries, J. C. Detter, E. Brambilla, M. Rohde, B. J. Tindall, M. Göker, T. Woyke, J. Bristow, J. A. Eisen, V. Markowitz, P. Hugenholtz, N. C. Kyrpides, H.-P. Klenk, A. Lapidus, Complete genome sequence of Cellulophaga algicola type strain (IC166T). *Stand. Genomic Sci.***4**, 72–80 (2011).
10. J. E. Lafleur, S. K. Costa, A. S. Bitzer, M. W. Silby, Draft Genome Sequence of Cellulophaga sp. E6, a Marine Algal Epibiont That Produces a Quorum-Sensing Inhibitory Compound Active against Pseudomonas aeruginosa. *Genome Announc.***3** (2015), doi:10.1128/genomeA.01565-14.
11. O. I. Nedashkovskaya, S. B. Kim, M. Vancanneyt, A. M. Lysenko, D. S. Shin, M. S. Park, K. H. Lee, W. J. Jung, N. I. Kalinovskaya, V. V. Mikhailov, K. S. Bae, J. Swings, Echinicola pacifica gen. nov., sp. nov., a novel flexibacterium isolated from the sea urchin Strongylocentrotus intermedius. *Int. J. Syst. Evol. Microbiol.***56**, 953–958 (2006).
12. H. Xu, Y. Fu, N. Yang, Z. Ding, Q. Lai, R. Zeng, Flammeovirga pacifica sp. nov., isolated from deep-sea sediment. *Int. J. Syst. Evol. Microbiol.***62**, 937–941 (2012).
13. W. Han, J. Gu, Q. Yan, J. Li, Z. Wu, Q. Gu, Y. Li, A polysaccharide-degrading marine bacterium Flammeovirga sp. MY04 and its extracellular agarase system. *J. Ocean Univ. China*. **11**, 375–382 (2012).
14. H. J. van der Meulen, W. Harder, H. Veldkamp, Isolation and characterization of Cytophaga flevensis sp. nov., a new agarolytic flexibacterium. *Antonie Van Leeuwenhoek*. **40**, 329–346 (1974).
15. A. J. Mann, R. L. Hahnke, S. Huang, J. Werner, P. Xing, T. Barbeyron, B. Huettel, K. Stüber, R. Reinhardt, J. Harder, F. O. Glöckner, R. I. Amann, H. Teeling, The Genome of the

Alga-Associated Marine Flavobacterium Formosa agariphila KMM 3901 T Reveals a Broad Potential for Degradation of Algal Polysaccharides. *Appl. Environ. Microbiol.***79**, 6813–6822 (2013).

16. R. Tanaka, Y. Mizutani, T. Shibata, H. Miyake, S. Iehata, T. Mori, K. Kuroda, M. Ueda, Genome Sequence of Formosa haliotis Strain MA1, a Brown Alga-Degrading Bacterium Isolated from the Gut of Abalone Haliotis gigantea. *Genome Announc.***4** (2016), doi:10.1128/genomeA.01312-16.
17. O. I. Nedashkovskaya, S. B. Kim, S. K. Han, A. M. Lysenko, M. Rohde, M.-S. Rhee, G. M. Frolova, E. Falsen, V. V. Mikhailov, K. S. Bae, Maribacter gen. nov., a new member of the family Flavobacteriaceae, isolated from marine habitats, containing the species Maribacter sedimenticola sp. nov., Maribacter aquivivus sp. nov., Maribacter orientalis sp. nov. and Maribacter ulvicola sp. nov. *Int. J. Syst. Evol. Microbiol.***54**, 1017–1023 (2004).
18. Y.-K. Kwon, J. H. Kim, J. J. Kim, S.-H. Yang, B.-R. Ye, S.-J. Heo, J.-H. Hyun, Z.-J. Qian, H.-S. Park, D.-H. Kang, C. Oh, Ochrovirga pacifica gen. nov., sp. nov., A Novel Agar-Lytic Marine Bacterium of the Family Flavobacteriaceae Isolated From A Seaweed. *Curr. Microbiol.***69**, 445–450 (2014).
19. W. Han, S. Zhao, H. Liu, Z. Wu, Q. Gu, Y. Li, [Isolation, identification and agarose degradation of a polysaccharide-degrading marine bacterium Persicobacter sp. JZB09]. *Wei Sheng Wu Xue Bao*. **52**, 776–83 (2012).
20. O. I. Nedashkovskaya, A. D. Kukhlevskiy, N. V. Zhukova, Polaribacter reichenbachii sp. nov.: A New Marine Bacterium Associated with the Green Alga Ulva fenestrata. *Curr. Microbiol.***66**, 16–21 (2013).
21. O. I. Nedashkovskaya, M. Suzuki, J.-S. Lee, K. C. Lee, L. S. Shevchenko, V. V. Mikhailov, Pseudozobellia thermophila gen. nov., sp. nov., a bacterium of the family Flavobacteriaceae, isolated from the green alga Ulva fenestrata. *Int. J. Syst. Evol. Microbiol.***59**, 806–810 (2009).
22. O. I. Nedashkovskaya, M. Suzuki, M. V. Vysotskii, V. V. Mikhailov, Reichenbachia agariperforans gen. nov., sp. nov., a novel marine bacterium in the phylum Cytophaga–Flavobacterium–Bacteroides. *Int. J. Syst. Evol. Microbiol.***53**, 81–85 (2003).
23. B. J. BACHMANN, Studies on Cytophaga fermentans, n.sp., a Facultatively Anaerobic Lower Myxobacterium. *J. Gen. Microbiol.***13**, 541–551 (1955).
24. S.-H. Yang, H.-S. Seo, J.-H. Woo, H.-M. Oh, H. Jang, J.-H. Lee, S.-J. Kim, K. K. Kwon, Carboxylicivirga gen. nov. in the family Marinilabiliaceae with two novel species, Carboxylicivirga mesophila sp. nov. and Carboxylicivirga taeanensis sp. nov., and reclassification of Cytophaga fermentans as Saccharicrinis fermentans gen. nov., comb. nov. *Int. J. Syst. Evol. Microbiol.***64**, 1351–1358 (2014).
25. J.-H. Yoon, S.-J. Kang, M.-H. Lee, T.-K. Oh, Tamlana agarivorans sp. nov., isolated from seawater off Jeju Island in Korea. *Int. J. Syst. Evol. Microbiol.***58**, 1892–1895 (2008).
26. R. Of, X. O. F. The, S12. t (2014).
27. M. JAM, D. FLAMENT, J. ALLOUCH, P. POTIN, L. THION, B. KLOAREG, M. CZJZEK, W. HELBERT, G. MICHEL, T. BARBEYRON, The endo-β-agarases AgaA and AgaB from the marine bacterium Zobellia galactanivorans: two paralogue enzymes with different molecular organizations and catalytic behaviours. *Biochem. J.***385**, 703–713 (2005).
28. L. Haridon, E. Corre, T. Barbeyron, B. Kloareg, P. Potin, Zobellia galactanovorans gen . nov ., sp . nov ., a marine species of Flavobacteriaceae isolated from a red alga , and classification of [ Cytophaga ] uliginosa ( ZoBell and Upham 1944 ) Reichenbach 1989 as Zobellia uliginosa gen .

nov ., comb . nov ., 985–997 (2001).

42. C. Oh, M. De Zoysa, Y.-K. Kwon, S.-J. Heo, A. Affan, W.-K. Jung, H.-S. Park, J. Lee, S.-K. Son, K.-T. Yoon, D.-H. Kang, Complete Genome Sequence of the Agarase-Producing Marine

Bacterium Strain S89, Representing a Novel Species of the Genus Alteromonas. *J. Bacteriol.* **193**, 5538–5538 (2011).
